# Warming During Embryogenesis Induces a Lasting Transcriptomic Signature in Fishes

**DOI:** 10.1101/2021.12.07.470713

**Authors:** Daniel M. Ripley, Terence Garner, Samantha A. Hook, Ana Veríssimo, Bianka Grunow, Timo Moritz, Peter Clayton, Holly A. Shiels, Adam Stevens

**Affiliations:** Division of Cardiovascular Sciences, Faculty of Biology, Medicine and Health, The University of Manchester, Manchester, UK; Division of Developmental Biology and Medicine, Faculty of Biology, Medicine, and Health, The University of Manchester, Manchester, UK; Department of Earth and Environmental Sciences, The University of Manchester, Manchester, UK; CIBIO, Centro de Investigação em Biodiversidade e Recursos Genéticos, InBIO Laboratório Associado, Campus de Vairão, Universidade do Porto, 4485-661 Vairão, Portugal; BIOPOLIS Program in Genomics, Biodiversity and Land Planning, CIBIO, Campus de Vairão, 4485- 661 Vairão, Portugal; Fish Growth Physiology, Research Institute for Farm Animal Biology (FBN), Wilhelm-Stahl-Allee 2, 18196 Dummerstorf, Germany; Deutsches Meeresmuseum, Katharinenberg 14–20, 18439 Stralsund, Germany; Institute of Biological Sciences, University of Rostock, Albert-Einstein-Straße 3, 18059 Rostock, Germany

**Keywords:** climate change, developmental plasticity, long-term effect, RNA-seq, network models

## Abstract

Exposure to elevated temperatures during embryogenesis can influence the plasticity of tissues in later-life. Despite these long-term changes in plasticity, few differentially expressed genes are ever identified, suggesting that the developmental programming of later-life plasticity may occur through the modulation of other aspects of the transcriptomic architecture, such as gene network function. Here, we use network modelling approaches to demonstrate that warm temperatures during embryonic development (developmental warming) have consistent effects in later-life on the organisation of transcriptomic networks across four diverse species of fishes: *Scyliorhinus canicula, Danio rerio, Dicentrarchus labrax*, and *Gasterosteus aculeatus*. The transcriptomes of developmentally warmed fishes are characterised by an increased entropy of their pairwise gene interaction networks, implying a less structured, more ‘random’ set of gene interactions. We also show that, in zebrafish subject to developmental warming, the entropy of an individual gene within a network is associated with that gene’s probability of expression change during temperature acclimation in later-life. However, this association is absent in animals reared under ‘control’ conditions. Thus, the thermal environment experienced during embryogenesis can alter transcriptomic organisation in later-life, and these changes may influence an individual’s responsiveness to future temperature challenges.

## Introduction

Developmental programming comprises changes to an organism’s phenotype that are induced by the conditions experienced during development (Bateson *et al*., 2014). These developmental conditions elicit changes in DNA and chromatin methylation patterns, causing alterations in gene expression (Anastasiadi *et al*., 2018, Anastasiadi *et al*., 2017, Fellous *et al*., 2015, Metzger and Schulte, 2017). Some of these changes in methylation pattern, and their concomitant influence on gene expression, can be retained into later-life, causing long-term physiological consequences for the individual. For instance, the developmental temperature of zebrafish (*Danio rerio*) affects their gene expression and physiology in adulthood, culminating in an enhanced thermal tolerance (Schaefer and Ryan, 2006) and acclimation capacity (Schnurr *et al*., 2014, Scott and Johnston, 2012). Studies in the threespined stickleback (*Gasterosteus aculeatus*) demonstrate that changes in developmental temperature cause DNA hypermethylation (Metzger and Schulte, 2017) and altered gene expression throughout adulthood (Metzger and Schulte, 2018). Thus, the embryonic environment can have long-term effects on an individual’s physiology.

As well as altering mean trait values, the embryonic environment can also affect an organism’s phenotypic plasticity, influencing its potential to acclimate to the conditions experienced in later-life (Beaman *et al*., 2016, Scott and Johnston, 2012, Loughland *et al*., 2021). That is, the capacity to physiologically respond to and mitigate future environmental challenges can be enhanced, hindered, or simply altered by an individual’s embryonic conditions (Beaman *et al*., 2016). Evidence from zebrafish (Scott and Johnston, 2012, Schnurr *et al*., 2014) and mosquitofish (*Gambusia holbrooki*) (Seebacher *et al*., 2014), amongst other species (Beaman *et al*., 2016), shows that the thermal environment experienced during early-life can have lasting effects on the acclimation capacity of multiple physiological traits, including metabolic rates (Seebacher *et al*., 2014), thermal tolerance (Healy *et al*., 2019), and swimming performance (Scott and Johnston, 2012).

Phenotypic changes arising from physiological plasticity are underwritten by changes in an organism’s gene expression in response to environmental cues. Such plasticity can be affected by an individual’s embryonic environment, which may either facilitate or hinder changes in later-life gene expression (Scott and Johnston, 2012, Beaman *et al*., 2016, Loughland *et al*., 2021). These changes in phenotypic plasticity, induced by developmental warming, are modulated through the activity of DNA methyltransferases (Loughland *et al*., 2021), which function to alter the methylation status of genes, influencing their expression. Thus, there is a link between an organism’s phenotype, gene expression, and plastic potential, which can be modulated by its embryonic environment (Beaman *et al*., 2016) via changes in DNA methylation (Loughland *et al*., 2021). Despite many recent studies demonstrating that the environmental conditions experienced during early-life often have lasting physiological consequences for an individual (Galli *et al*., 2021, Ruhr *et al*., 2021, Hellgren *et al*., 2021, Ruhr *et al*., 2019, Seebacher *et al*., 2014, Scott and Johnston, 2012), including changes in phenotypic plasticity (Scott and Johnston, 2012), very few developmentally programmed, differentially expressed genes (DEGs) are ever identified.

Recent advances in gene expression analyses demonstrate that transcriptomic mechanisms aside from the differential expression of genes can contribute to an organism’s physiology and physiological plasticity. Network analyses, rather than focussing on quantifying changes in gene expression, assess the role of genes within an expression network (Torson *et al*., 2020, Banerj *et al*., 2013). By assessing genes within networks, rather than as isolated expression counts, differences in organismal physiology can be associated with changes in gene network function, regardless of whether individual genes show differential expression. For example, gene co-expression networks have been used successfully in mammals to show that *NUEROD1* and *RCAN1*, genes known to be associated with diabetes (Malecki *et al*., 1999, Peiris *et al*., 2012), are not highlighted as interesting by differential expression analysis between diabetic and control groups, but are physiologically important owing to their altered function within transcriptional networks (Iacono *et al*., 2019). In the southern grey-cheeked salamander (*Plethodon metcalfi*), transcriptional network analyses have implicated blood vessel morphogenesis in the plasticity of desiccation tolerance (Riddell *et al*., 2019). Therefore, by assessing transcriptional networks, novel insight into physiological phenomena can be gained that differential expression analyses alone cannot capture (Iacono *et al*., 2019, Torson *et al*., 2020).

Studies analysing gene expression networks typically represent gene-gene interactions as a grid (matrix) of pairwise relationships. Pairwise networks use gene expression count data to define gene-gene interactions based on correlations between the expression of pairs of genes. Pairwise networks capture interesting and relevant biological information beyond that which can be identified through differential gene expression analysis alone (Iacono *et al*., 2019, Riddell *et al*., 2019). However, these simple pairwise networks do not capture the nuance of higher order interactions that can be crucial to the understanding of biological systems (Murgas *et al*., 2022, Battiston *et al*., 2020, Mickalide and Kuehn, 2019, Sanchez, 2019). Higher order interactions are correlations that are modified, or only exist, when three or more components are present (Sanchez, 2019). Rather than modelling direct connections, higher order interaction networks cluster genes based on their shared connections to the rest of the transcriptome. Thus, higher order interaction networks allow more distant relationships within the transcriptome to be modelled. Previous studies have incorporated higher order interactions into gene co-expression network analyses (e.g. Ruane *et al*., 2022).

Various properties of both pairwise and higher order gene expression networks can be described using mathematical tools from the field of graph theory (Murgas *et al*., 2022, Banerj *et al*., 2013, Teschendorff and Enver, 2017, Iacono *et al*., 2019). Shannon entropy (herein entropy) has shown particular promise for capturing biological information relevant to plasticity at the cellular level (Banerj *et al*., 2013, Teschendorff and Enver, 2017). Transcriptional network entropy is a metric describing the structure of gene interaction networks (see supplementary Figure 1 for a conceptual overview). Network entropy is known to correlate with the differentiation potential and plasticity of cells (Teschendorff and Enver, 2017, Banerji *et al*., 2013). As a cell becomes more differentiated, its signalling promiscuity (entropy) decreases as the gene-gene interactions become more specialised and thus less random (Banerji *et al*., 2013, Teschendorff and Enver, 2017). Similarly, upon exposure to a stressor, gene expression entropy decreases, likely due to the activation of highly coordinated, stressor-specific pathways (Ogata *et al*., 2015). Indeed, the ability to produce a highly coordinated response to environmental challenges is a strong predictor of survival under extreme stress (Zhu *et al*., 2020). Thus, transcriptomic entropy is a useful metric of gene expression coordination (Banerji *et al*., 2013, Teschendorff and Enver, 2017) that is known to change following environmental stress (Zhu *et al*., 2020), likely due to the coordination of a response.

Given that transcriptomic entropy is associated with both organismal responses to environmental stress (Zhu *et al*., 2020) and the plasticity of cells (Teschendorff and Enver, 2017, Banerj *et al*., 2013, Park *et al*., 2016), we reasoned that entropy may convey information pertinent to the plasticity of groups of cells (i.e. tissues) during environmental changes too. The physiological plasticity of ectotherms is known to be affected by the embryonic environment (Scott and Johnston, 2012, Seebacher *et al*., 2014, Beaman *et al*., 2016) via changes in methylation (Loughland *et al*., 2021). Despite identifying changes in DNA methylation status (Ruhr *et al*., 2021, Metzger and Schulte, 2017) and organismal physiology (Galli *et al*., 2023), very few studies of ectotherm developmental programming observe substantial differences in gene expression (Scott and Johnston, 2012, Metzger and Schulte, 2018, Anastasiadi *et al*., 2017), suggesting that the physiological effects associated with the changes in DNA methylation may occur through the modulation of other aspects of the transcriptomic architecture, such as network function. Consequently, we hypothesise that warming during embryogenesis alters the organisation of transcriptional networks, and that these changes in network organisation influence the plasticity of gene expression in later-life.

Here, we test the effects embryonic temperature has on the entropy of transcriptomic networks in four species of fishes: the small spotted catshark (*Scyliorhinus canicula*, ventricular tissue), zebrafish (*Danio rerio*, fast hypaxial muscle), European seabass (*Dicentrarchus labrax*, muscle), and threespine stickleback (*Gasterosteus aculeatus*, muscle), before assessing whether any changes in network organisation, caused by the developmental environment, are associated with an increased transcriptomic sensitivity to future thermal acclimation.

## Materials and Methods

### Experimental Animals

Details of the experimental design can be found in Figure 1. The raw *D. rerio, D.labrax*, and *G. aculeatus* data are published in Scott and Johnston 2012, Anastasiadi *et al*., 2021, and Metzger and Schulte 2018, respectively. The *S. canicula* data were collected *de novo* for this study.

**Figure 1:**
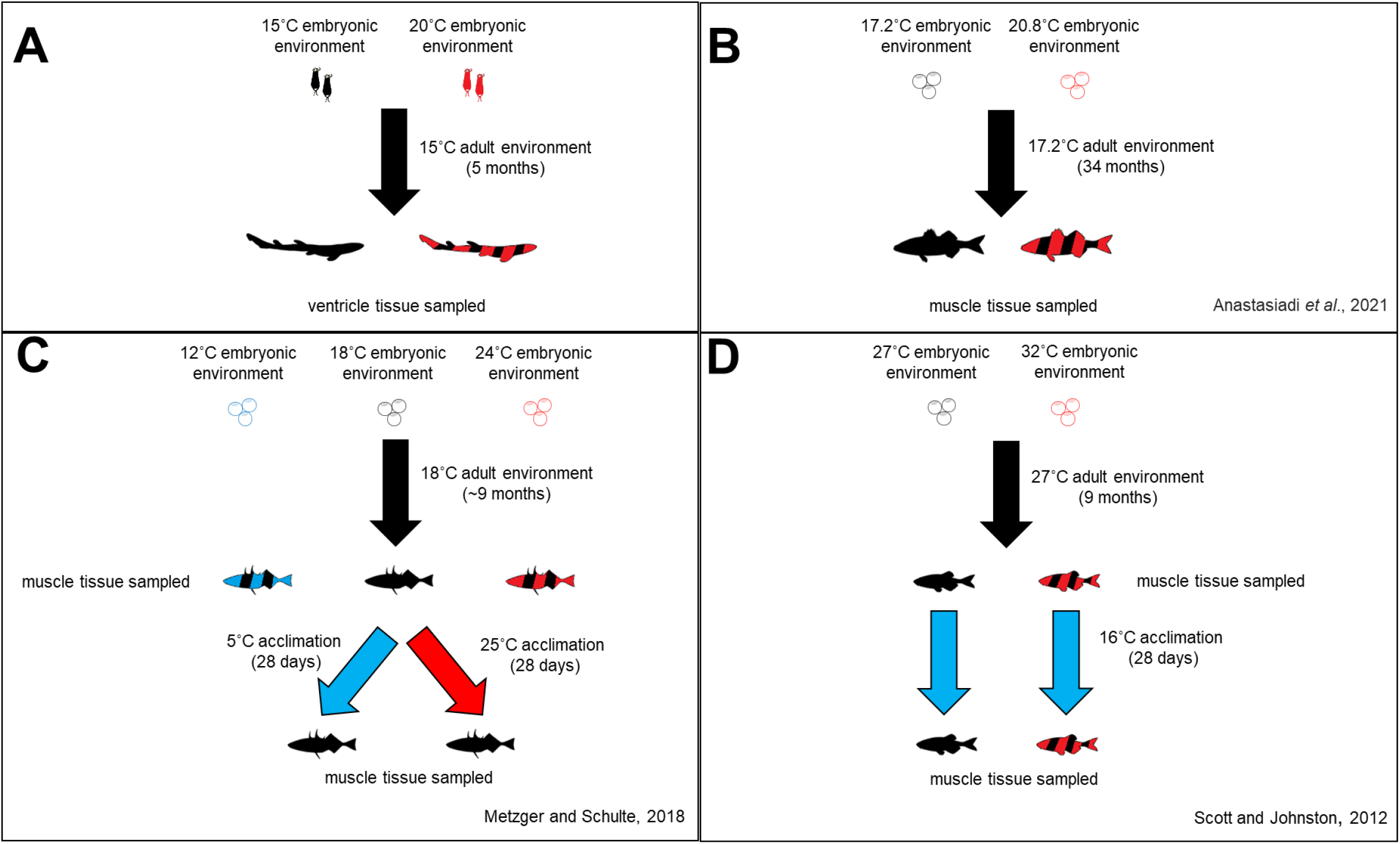
An overview of the experimental design for the RNA-seq data collected from **A)** *Scyliorhinus canicula*, and re-analysed from **B)** *Dicentrarchus labrax* (Anastasiadi *et al*., 2021), **C)** *Gasterosteus aculeatus* (Metzger and Schulte, 2018), and **D)** *Danio rerio* (Scott and Johnston, 2012).

Small-spotted catshark embryos were collected from a population of 7 randomly mating adult individuals held at 15 ºC at the Ozeaneum, Stralsund, Germany, and transported to the University of Manchester, UK. Upon arrival the health and developmental stage of each embryo was assessed using the Musa scale (Musa *et al*., 2018). Only healthy embryos at stage 1 were used in the study (Musa *et al*., 2018). All regulated procedures received approval from the institution’s ethical review board and were performed under the Home Office License P005EFE9F9 held by HAS.

Following embryonic staging, individuals were randomly assigned to a temperature treatment group (15 ± 0.3 ºC or 20 ± 0.3 ºC). For the control group, 15 ºC was chosen as it falls within the range of temperatures experienced by *S. canicula* in the wild and is the holding temperature of the parent population at the Ozeaneum, Stralsund, Germany. For the treatment group, +5 ºC (20 ºC) was chosen as it represents the increase in ocean temperature predicted by the end of the century, whilst also being within the current range of temperatures experienced by some *S. canicula* populations in the wild (Pegado *et al*., 2020). The egg cases in each treatment group were hung vertically in two, well aerated, 55 l static seawater (35 ppt salinity) tanks equipped with internal filters and left to continue their embryonic development.

Upon hatching, fin clips were taken to facilitate identification in later-life through microsatellite analysis. The fin clips were stored in 98% ethanol at -20 ºC, and the sharks were moved into one of four well aerated 400 l static seawater tanks held at 15 ºC ± 0.3 ºC. The hatchlings from the 20 ºC treatment group were lowered to 15 ºC at a rate of 2.5 ºC per day, and sharks from both treatment groups were mixed and randomly allocated to a tank. The sharks were fed a mixture of squid, crab and krill three times per week. The sharks from both treatment groups were held at 15 ºC ± 0.3 ºC for 4-5 months prior to tissue sampling. During the entire experiment, saltwater changes were performed three times weekly to maintain ammonia, nitrite, and nitrate below detectable levels.

### Tissue Sampling

Sharks at 4-5 months age (mean ± SEM = 136 ± 6.3 days) were euthanised with an overdose of buffered tricane methanesulfonate and a fin clip was taken post-humous. Fin clips were snap frozen in liquid nitrogen and stored at -80 ºC. The ventricle was excised from each individual and frozen in liquid nitrogen before being stored at -80 ºC prior to RNA-sequencing (RNA-seq). Ventricle tissue was chosen for sequencing owing to the ventricle’s key role in mediating thermal performance and tolerance (Pörtner and Knust, 2007). Fin clips were also taken from anaesthetised animals in the parent population and stored at -80 ºC.

### Individual Identification Using Microsatellite Analysis

To identify which condition an individual was developed in, microsatellite analysis was performed on the fin clips taken at birth and at the end of the experiment to match each individual using the methods described in Hook *et al*. (2019a) and Hook *et al*. (2019b) Fin clips of the parents were also extracted and genotyped using the same method. Fin clips were extracted using a BioLine Isolate Genomic kit with an extended proteinase K digestion to maximise DNA yield. DNA yield was assessed using a NanoDrop ND-1000 spectrophotometer (Thermo Fisher Scientific, USA) and gel electrophoresis. A one-primer cocktail containing 11 microsatellites and three tail dyes was used for DNA amplification (Griffiths *et al*., 2011) using the QIAGEN multiplex PCR kit. The thermocycling protocol consisted of initial denaturation cycle at 95 ºC for 15 minutes, followed by 35 cycles of 94 ºC for 30 seconds, 60 ºC for 90 seconds and 72 ºC for 45 seconds, and finalised by one cycle at 72 ºC for 30 minutes. The products were visualised on a gel and then genotyped using an ABI sequencer. Positive control samples were added to each plate genotyped to account for possible allele slippage.

Genotypes were scored using GeneMapper© v4.1 (Applied Biosystems) and validated through Microchecker (van Oosterhout *et al*., 2004). Duplicates were found between the two batches using CERVUS (Marshall *et al*., 1998). In cases where some alleles within the genotype did not match, controls were used as a reference to identify possible allele slippage. Probabilities of identity analysis were taken from CERVUS to confirm match identification between the two batches and the parentage of each individual. This analysis allowed us to identify the developmental group each shark came from whilst minimising potential biases by blinding the experimenters to the individual’s developmental conditions until after the ventricle was dissected. The probability of identity analysis also allowed us to confirm that no samples sent for RNA-seq came from full siblings.

### *de-novo* Transcriptome Assembly and Differential Expression Analysis

The ventricle tissue from six individuals (N = 3 per treatment group), stored in RNAlater at -80 ºC, was used for generating the transcriptome. The tissue samples were extracted using the Qiagen kit with homogenisation. Total RNA was submitted to the Genomic Technologies Core Facility (GTCF) at The University of Manchester. Quality and integrity of the RNA samples were assessed using a 2200 TapeStation (Agilent Technologies). Libraries were generated using the TruSeq® Stranded mRNA assay (Illumina, Inc.) according to the manufacturer’s protocol. mRNA enrichment was performed using poly adenosine selection. 76 base pair long, pair ended sequencing was performed using an Illumina HiSeq4000. Reads that mapped to human, bacterial, and viral sequences were removed using DeconSeq (Schmieder and Edwards, 2011) whilst reads mapping to ribosomal RNA were removed using sortmeRNA (Kopylova *et al*., 2012). Sequences with a low-quality score (regions averaging a score <5 over a 4bp sliding window, and leading/trailing sequences scoring <5) were then removed from the cleaned reads using Trimmomatic (Bolger *et al*., 2014) prior to transcriptome assembly. 264 million bases passed the quality control process and were used for transcriptome assembly in Trinity 2.2.0 (Grabherr *et al*., 2011, Haas *et al*., 2013). Default parameters were used in Trinity, except –normalize_reads, producing 347710 contigs with a N50 of 1422, and a BUSCO score of 88.8%, with 519 single copy, and 2458 duplicated copy, BUSCOs (BUSCO v2.2, Simao *et al*., 2015). Open reading frames (ORFs) were predicted from the transcripts using Transdecoder (https://github.com/TransDecoder/TransDecoder/wiki) with a minimum length threshold of 100 amino acids. This filtered dataset was then annotated using a BLAST search against the SwissProt database. The reads were pseudo-aligned to the curated transcript assemblies using Kallisto (Bray *et al*., 2016), allowing the relative abundance of each transcript to be calculated. Where multiple transcripts mapped to a single BLAST hit, the sum of the transcript abundances was used. The abundance estimates were filtered by removing fragments that had ≤ 0.5 counts per million before differential expression analysis was performed with EdgeR (McCarthy *et al*., 2012, Robinson *et al*., 2010) in R 4.1.2 (R Core Team, 2019) using the R-script available in Metzger and Schulte (2018). Significance levels were adjusted for multiple testing using a false discovery rate (FDR) correction.

### Entropy Analysis

Gene networks, utilising either pairwise or higher order interactions, were generated for each treatment group. The expression of each gene was tested for correlations against all the assembled transcripts within each treatment group. The correlation values were binarized such that strong positive (top 10% of r values, *S. canicula* r ≥ 0.95, *D. rerio* r ≥ 0.81, *D. labrax* ≥ 0.72, *G.aculeatus* r ≥ 0.62) and strong negative (bottom 10% of r values, *S. canicula* r ≤ -0.95, *D. rerio* r ≤ -0.80, *D. labrax* ≤ -0.70, *G.aculeatus* r ≤ -0.61) correlations were retained (denoted as 1) and weak correlations were discarded (denoted as 0). This binarized correlation matrix (*M*) is square and represents the pairwise interaction network.

To generate the higher order interaction networks (hypernetworks), the binarized correlation matrix was multiplied by the transpose of itself (*M* × *M*^*T*^) to return the adjacency matrix of the hypernetwork, where the values in each cell represent the number of shared correlations any given pair of genes has to the rest of the transcriptome. Thus, the higher order network models relationships between genes, which neither rely on, nor are predicated solely on, the existence of a pairwise correlation between those genes.

To assess properties of the whole-transcriptome, subsets of 200 genes were randomly selected (with replacement) from each network and the entropy of each gene within the subset was calculated using the R packages mixOmics (Rohart *et al*., 2017) and BioQC (Zhang *et al*., 2017) in R 4.1.2 (R Core Team, 2019). The entropy (Shannon, 1948) of a gene was defined as

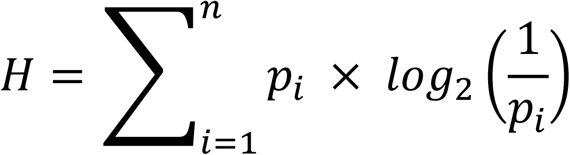

where *H* is entropy, and *p* is the probability of outcome *i*.

High entropy hypernetworks comprise genes with ‘random’ numbers of shared correlations. Conversely, low entropy networks contain peaks and troughs in the probability distribution, such that certain numbers of shared correlations are more likely than others. Thus, movement from high entropy (random) to low entropy (structured) networks conveys information about increasing transcriptomic structure. This approach has previously been applied to transcriptomic datasets (e.g. Ruane *et al*., 2022).

The mean entropy was calculated for each subset of genes (n = 200 genes). This approach was iterated 1000 times to provide a whole-transcriptome assessment of both the pairwise and higher order (hypernetwork) network entropy for animals from each treatment group. This iterative approach using sub-networks of a uniform size was required as entropy values are sensitive to the size of the transcriptome, which differs subtly between treatment groups. Wilcoxon tests were used to contrast network entropies between the fish from different developmental backgrounds. P-values were adjusted for multiple comparisons testing using the Benjamini-Hochberg method.

As bootstrapping (1000 iterations) was used to generate the entropy data, we performed a supplementary analysis that is less sensitive to n-number to ensure that any significant differences identified between the treatment groups were not artefacts of the high n-number. To do this, we created logistic regression models (R package ‘lme4’ (Bates *et al*., 2015)) for each species with developmental group as the response variable and entropy as the predictor. From each model we calculated concordance (C), a value between 0 and 1 that represents the discriminatory ability of a model. A concordance value of 0.5 represents a model with no predictive power (i.e. random guessing) whilst a concordance of 1.0 shows a model that can discriminate between two groups with 100% accuracy. Concordance values and their associated 95% confidence intervals were calculated.

### *Danio rerio, Dicentrarchus labrax*, and *Gasterosteus aculeatus* Re-analysis

The computational analysis pipeline described above was also implemented on previously published RNA-seq datasets of temperature-induced developmental programming in zebrafish (*D. rerio*) hypaxial fast muscle (Scott and Johnston, 2012), European seabass (*D. labrax*) muscle tissue (Anastasiadi *et al*., 2021), and threespine stickleback (*G. aculeatus*) muscle tissue (Metzger and Schulte, 2018). The *D. rerio, D. labrax*, and *G. aculeatus* data were processed in an identical manner to the *S. canicula* data. Developmental conditions and transcriptome assembly statistics for the four species are shown in Table 1. Full details of the animals used in the re-analysis can be found in Scott and Johnston 2012 (*D. rerio*, N = 4), Anastasiadi *et al*. 2021 (*D. labrax*, N = 5), and Metzger and Schulte 2018 (*G. aculeatus*, N = 6), respectively.

**Table 1.** Experimental details and transcriptome assembly statistics for *Scyliorhinus canicula, Danio rerio* (Scott and Johnston 2012), *Dicentrarchus labrax* (Anastasiadi *et al*., 2021) and *Gasterosteus aculeatus* (Metzger and Schulte, 2018).

| Parameter | Species |  |  |  |
| --- | --- | --- | --- | --- |
|  | *Scyliorhinus canicula* | *Danio rerio* | *Dicentrarchus labrax* | *Gasterosteus aculeatus* |
| time since exposure | 4-5 months | 8-9 months | 34 months | ~9 months |
| developmental temperatures | 15°C and 20°C | 27°C and 32°C | 17.2°C and 20.8°C | 12°C, 18°C and 24°C |
| later-life acclimation temperature(s) | 15°C | 27°C and 16°C | 17.2°C | 18°C, 25°C, and 5°C |
| exposure period | until hatching (~15 weeks) | until hatching (~3 days) | 6-63dpf (57 days) | until hatching (5-23 days) |
| life stage at sampling | juvenile | adult | adult | adult |
| tissue sampled | ventricle | hypaxial fast muscle | muscle | muscle |
| N50 | 1422 | 916 | 1601 | 1743 |
| median contig length | 363 | 325 | 428 | 425 |

### Acclimation Analysis

Only the zebrafish (*D. rerio*) and stickleback (*G*. *aculeatus*) were acclimated to different temperatures in adulthood (see Figure 1). Therefore, the acclimation analysis is limited to these two species.

The acclimation analysis allowed us to test two hypotheses: 1) that a gene’s entropy within expression networks would be associated with the probability of its expression changing following thermal acclimation; 2) that developmental temperature would interact with a gene’s entropy to influence the likelihood of its expression changing following thermal acclimation.

### Acclimation Analysis: *Danio rerio*

A subset of zebrafish from both developmental temperatures (27 ºC and 32 ºC) were cold acclimated (16 ºC for 28-30 days, N = 4) in adulthood prior to RNA-sequencing (Scott and Johnston 2012). We re-analysed these data to investigate whether the whether transcriptomic responsiveness to cold acclimation in adulthood differed between developmental groups, and if any differences were associated with the organisation of the gene networks (assessed via entropy).

The proportion of genes whose expression changed due to cold acclimation was compared using a Chi squared test with Yates’ continuity correction. The magnitude of change in the DEGs was contrasted by taking the modulus of each gene’s log fold change and comparing it between groups using a Wilcoxon test. These analyses were performed to confirm that our RNA-seq re-analysis matched the original findings of Scott and Johnston (2012).

To build on this confirmatory analyses and incorporate the data into the framework of network organisation, we estimated the probability of gene expression change using a logistic regression model (R package ‘lme4’ (Bates *et al*., 2015)) to assess whether a gene’s response (i.e. differentially expressed or not, as described previously) to later-life cold-acclimation was influenced by its pre-acclimation entropy. Separate models were created to assess the influence of a gene’s pairwise and higher order network entropy on the probability of its expression changing following future cold acclimation. Entropy (either pairwise or higher order), developmental temperature, and their interaction were included as predictors in the model. Significance was assessed using a Wald test (package ‘aod’, Lesnoff and Lancelot, 2012).

### Acclimation Analysis: *Gasterosteus aculeatus*

A subset of stickleback from the control developmental temperatures only (18 ºC) were cold (5 °C for 4 weeks, N = 6) or warm (25 °C for 4 weeks, N = 6) acclimated in adulthood prior to RNA-sequencing (Metzger and Schulte 2018). These data allowed us to assess whether transcriptomic organisation (as defined by entropy) prior to later-life temperature acclimation was associated with the likelihood of a gene’s expression changing following cold or warm acclimation. To assess this, a logistic regression model (R package ‘lme4’ (Bates *et al*., 2015)) was created to determine whether a gene’s response to temperature acclimation in adulthood (as defined by a false discovery rate corrected p-value (FDR) cut-off of 0.05) was associated with its entropy prior to the later-life thermal acclimation. Separate models were created to assess the influence of a gene’s pairwise and higher order network entropy on the probability its expression changing following future thermal acclimation. Entropy (either pairwise or higher order), acclimation temperature (5 °C or 25 °C), and their interaction were included as predictors in the model. Significance was assessed using a Wald test (package ‘aod’, Lesnoff and Lancelot, 2012). This analysis could only be performed on stickleback from a control embryonic background.

## Results

### Differential Gene Expression Analysis

Developmental warming caused the differential expression of 164 (FDR < 0.05) genes in juvenile *S. canicula*, compared to 64 (FDR < 0.05), 79 (FDR < 0.05), and 68 (FDR < 0.05) in adult *D. labrax, D. rerio*, and *G. aculeatus* respectively (Figure 2). The DEGs showed little overlap between species (Figure 2).

**Figure 2:**
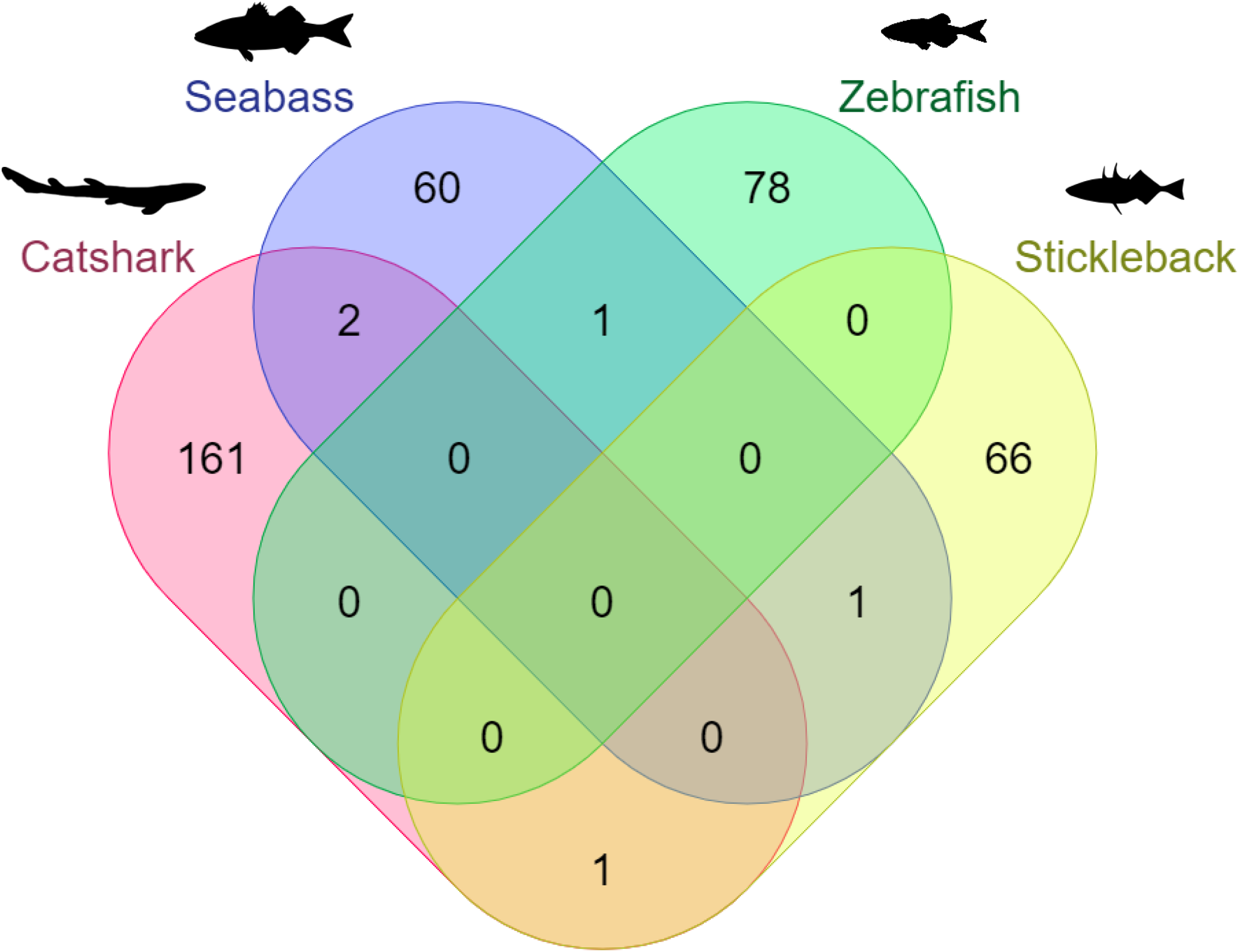
Overlap of the genes in *Scyliorhinus canicula* (red), *Dicentrarchus labrax* (blue), *Danio rerio* (green), and *Gasterosteus aculeatus* (yellow) that are differentially expressed (FDR < 0.05) in later-life due to increased embryonic temperature. *D. labrax, D. rerio, and G. aculeatus* data are re-analysed from Anastasiadi *et al*., 2021, Scott and Johnston 2012, and Metzger and Schulte 2018, respectively.

### Pairwise Network Entropy Analysis

Entropy of the pairwise interaction networks was increased by developmental warming in *S. canicula* (Figure 3A, p < 0.0001, C ± 95% CI = 0.90 ± 0.014), *D.labrax* (Figure 3B, p < 0.0001, C ± 95% CI = 0.87 ± 0.016), *D. rerio* (Figure 3C, p < 0.0001, C ± 95% CI = 1.00 ± 0.00), and *G. aculeatus* (Figure 3D, p < 0.0001, C ± 95% CI = 0.82 ± 0.016). In *G. aculeatus*, developmental cooling reduced the entropy of the pairwise interaction networks (Figure 3D, p < 0.0001, C ± 95% = 0.82 ± 0.016). The increases in transcriptomic network entropy caused by developmental warming demonstrate differences in the ‘randomness’ of the pairwise interaction networks. Thus, these data suggest that developmental warming results in less structured (more random) gene interactions in four species of fishes: *S. canicula, D. labrax, D. rerio*, and *G. aculeatus*.

**Figure 3:**
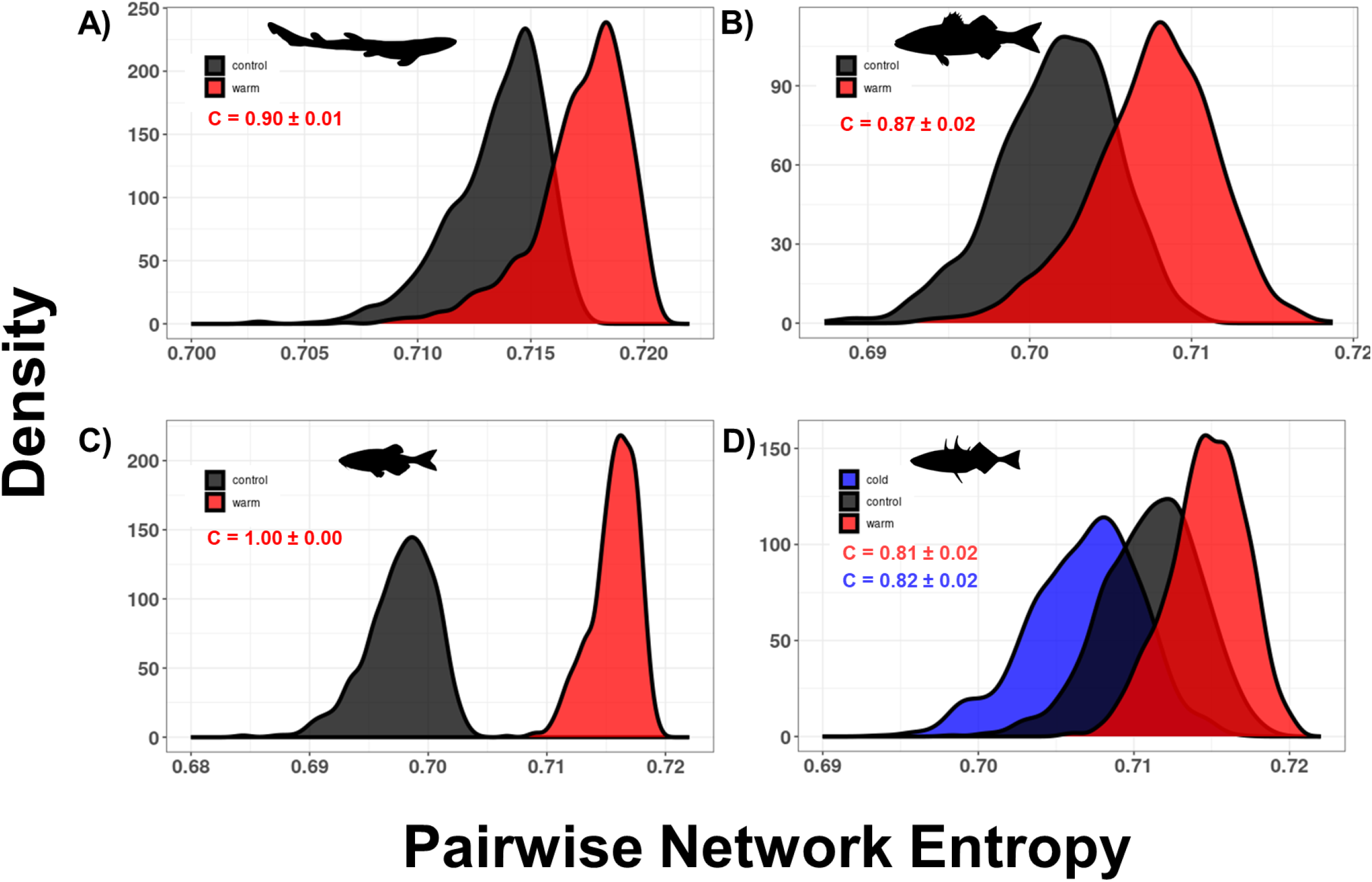
Density plots displaying the distribution of mean entropy values from the **pairwise gene interaction networks** in **A)** *Scyliorhinus canicula*, **B)** *Dicentrarchus labrax*, **C)** *Danio rerio*, and **D)** *Gasterosteus aculeatus* that underwent embryogenesis at different temperatures. Inset shows concordance ± 95% confidence intervals. *Scyliorhinus canicula*: n = 1000 iterated gene networks, N = 3 animals per group. *Dicentrarchus labrax*: n = 1000 iterated gene networks, N = 5 animals per group. *Danio rerio*: n = 1000 iterated gene networks, N = 4 animals per group. *Gasterosteus aculeatus*: n = 1000 iterated gene networks, N = 6 animals per group. *D. labrax, D. rerio, and G. aculeatus* data are re-analysed from Anastasiadi *et al*., 2021, Scott and Johnston 2012, and Metzger and Schulte 2018 respectively.

### Hypernetwork Entropy Analysis

Entropy of the higher-order interaction networks was also increased by developmental warming in *S. canicula* (Figure 4A, p < 0.0001, C ± 95% CI = 0.59 ± 0.026), *D.labrax* (Figure 4B, p < 0.0001, C ± 95% CI = 0.89 ± 0.014) and *D. rerio* (Figure 4C, p < 0.0001, C ± 95% CI = 1.00 ± 0.00), but not in *G. aculeatus*, where entropy of the higher-order interactions was highest in the control animals, and reduced slightly by both developmental warming and cooling (Figure 4D, p < 0.0001, C ± 95% CI = 0.55 ± 0.022 and p < 0.0001, C ± 95% CI = 0.59 ± 0.024). It is important to note the low concordance values for the stickleback hypernetwork data, indicating that whilst the p-values show significance, there is considerable overlap in entropy values between the different developmental cohorts.

**Figure 4:**
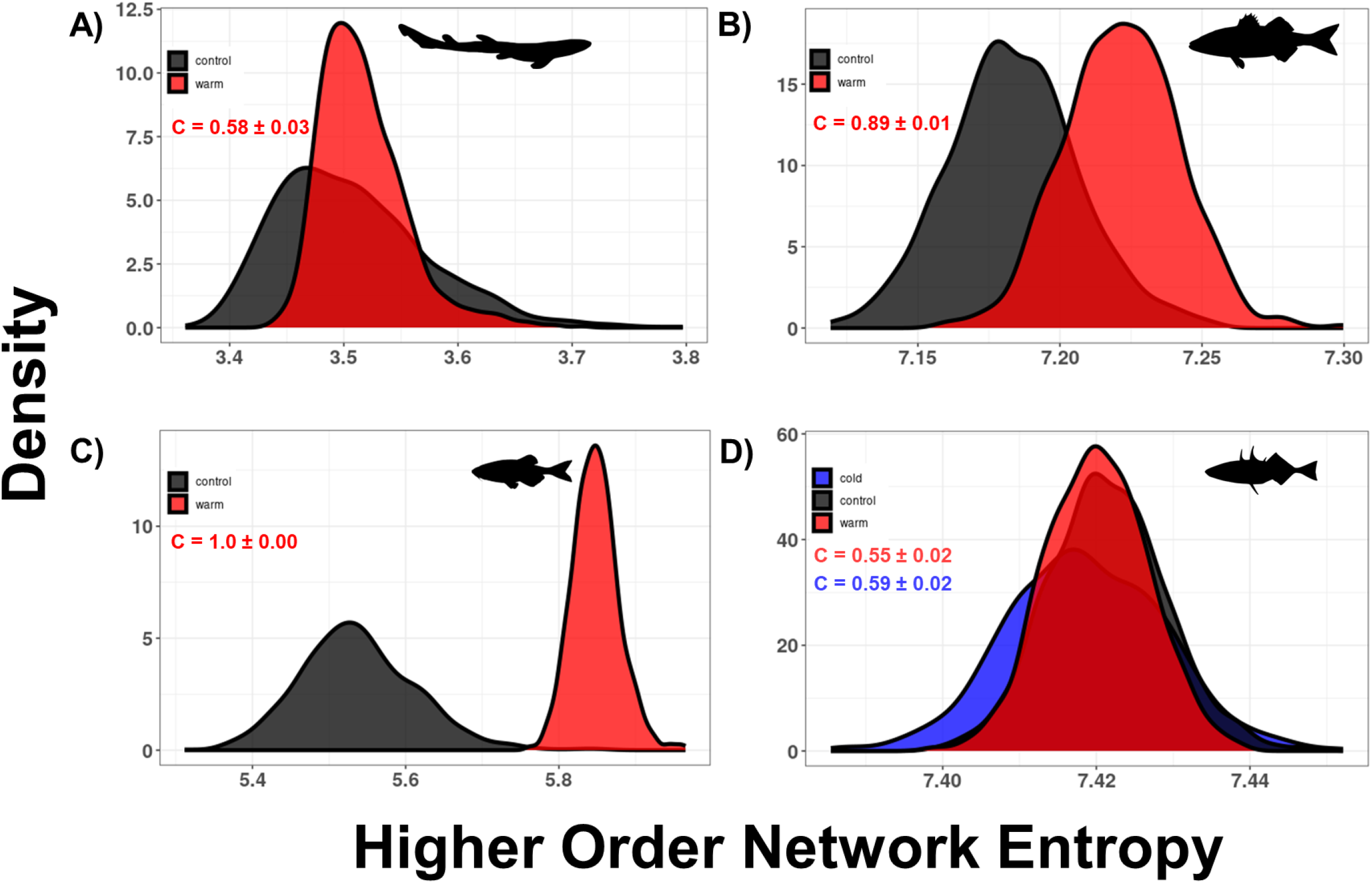
Density plots displaying the distribution of mean entropy values from the **higher order gene interaction networks** in **A)** *Scyliorhinus canicula*, **B)** *Dicentrarchus labrax*, **C)** *Danio rerio*, and **D)** *Gasterosteus aculeatus* that underwent embryogenesis at different temperatures. Inset shows concordance ± 95% confidence intervals. *Scyliorhinus canicula*: n = 1000 iterated gene networks, N = 3 animals per group. *Dicentrarchus labrax*: n = 1000 iterated gene networks, N = 5 animals per group. *Danio rerio*: n = 1000 iterated gene networks, N = 4 animals per group. *Gasterosteus aculeatus*: n = 1000 iterated gene networks, N = 6 animals per group. *D. labrax, D. rerio, and G. aculeatus* data are re-analysed from Anastasiadi *et al*., 2021, Scott and Johnston 2012, and Metzger and Schulte 2018 respectively.

These data suggest that developmental warming results in less structured (more random) higher order gene interactions of *S. canicula, D. labrax*, and *D. rerio*, as was seen in the pairwise network analysis. However, this is not true of *G. aculeatus*.

### Acclimation Analysis

#### Danio rerio

*D. rerio* that were reared at an elevated temperature showed a greater number (p < 0.0001) and magnitude (p < 0.0001) of gene expression changes following 28-30 days cold acclimation (16 ºC) than *D. rerio* developed in control conditions. These analyses simply confirm that our transcriptome assembly and DEG analysis match the original findings of Scott and Johnston (2012).

Logistic regression models were used to assess whether the entropy of the pairwise (McFadden’s pseudo-r^2^ = 0.0037) and higher order (McFadden’s pseudo-r^2^ = 0.0032) interaction networks influenced the response of a fish’s transcriptome to a later-life thermal challenge (16 ºC acclimation, see Figure 1).

Developmental group (p = 0.015), and the interaction between developmental group and pairwise network entropy (p = 0.0033), affected gene expression change following a thermal challenge at the adult stage (Figure 5c). Pairwise network entropy alone had no significant effect (p = 0.42).

**Figure 5:**
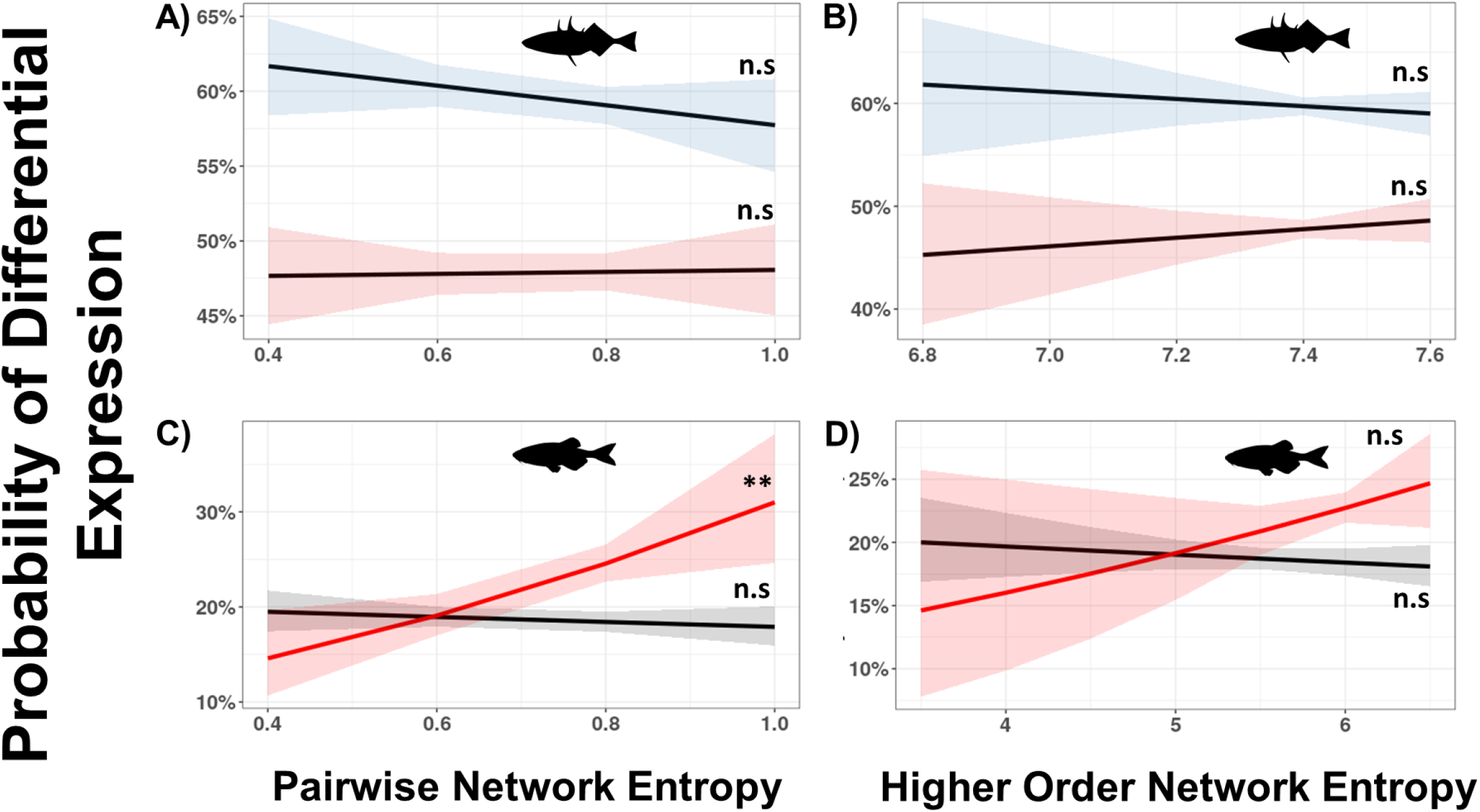
Graphs showing the relationship between a gene’s entropy and the probability of its expression changing following thermal acclimation. Panels **A)** and **B)** show *Gasterosteus aculeatus* incubated in control conditions throughout embryogenesis and acclimated to warm (red) or cold (blue) conditions in adulthood. Panels **C)** and **D)** show *Danio rerio* incubated in control (black) or warm (red) treatments throughout embryogenesis, raised in control conditions until adulthood, and then acclimated to cold temperatures. ** represents a relationship between entropy and the probability of gene expression following acclimation (p < 0.01) n = 13441 and 8614 for genes within the *Gasterosteus aculeatus* and *Danio rerio* transcriptomes respectively. N = 6 and 4 animals used to generate the gene expression networks for *Gasterosteus aculeatus* and *Danio rerio* respectively. Data are reanalysed from Metzger and Schulte 2018 and Scott and Johnston 2012.

No significant effect of higher order network entropy (p = 0.41), or its interaction with developmental temperature (p = 0.11), was observed (Figure 5D).

#### Gasterosteus aculeatus

Only stickleback from the ‘control’ developmental background were warm or cold acclimated during later-life (Metzger and Schulte, 2018). Thus, the stickleback re-analysis could be used only to test for an association between a gene’s entropy and its probability of differential expression following later-life thermal acclimation in fish from control conditions.

There was no association between a gene’s pairwise (Figure 5A, p = 0.21) or higher order (Figure 5B, p = 0.53) network entropy and its probability of expression following later-life thermal acclimation. No significant interactions between acclimation temperature and pairwise (Figure 5A, p = 0.32) or higher order (Figure 5B, p = 0.33) network entropy were observed (pairwise model, McFadden’s pseudo-r^2^ = 0.019; higher order model, McFadden’s pseudo-r^2^ = 0.019).

## Discussion

Previous studies have shown that the thermal environment experienced during embryogenesis can have persistent effects on the physiology and plasticity of muscle tissue in fishes (Schnurr *et al*., 2014, Scott and Johnston, 2012, Beaman *et al*., 2016). Here, we show that despite causing few and dissimilar DEGs, elevated developmental temperature induces consistent changes in the organisation of transcriptional networks across four species of fish. Furthermore, in zebrafish, we demonstrate that developmental warming results in a positive association between a gene’s entropy and the probability of its expression changing following a thermal challenge at the adult stage. Thus, these data suggest that by altering the organisation of transcriptomic networks, warming experienced during embryogenesis may influence the response of a tissue’s transcriptome to thermal challenges throughout the individual’s lifetime. Interestingly, animals from a ‘control’ developmental background showed no relationship between transcriptomic entropy and differential expression following thermal acclimation.

Consistent with previous studies, we found that developmental warming affected the expression of few (68-164) genes in later-life, with little overlap between species (Figure 2, Scott and Johnston, 2012, Metzger and Schulte, 2018, Anastasiadi *et al*., 2021). Despite little overlap in the DEGs, a consistent increase in the entropy of pairwise gene interaction networks was observed with increased developmental temperature in each species (Figure 3). Whilst the n-numbers per species are modest (3 – 6 fish per treatment group per species), finding a consistent pattern across four species lends weight to this observation. Entropy of the higher order interaction networks was also affected by developmental warming, increasing in the catshark, seabass, and zebrafish, but decreasing subtly in the stickleback (Figure 4). Whilst it is tempting to speculate on the reason that the higher order interaction networks of the stickleback differ to those of the catshark, zebrafish, and seabass, especially given the stickleback’s complex evolutionary history (McKinnon and Rundle, 2002), the data required to disentangle this question do not yet exist.

Consistent differences in the organisation of pairwise networks across species suggests that the embryonic environment may influence the function of gene networks in general (Banerj *et al*., 2013, Park *et al*., 2016). Indeed, we identify a common transcriptomic response to early-life warming across four species of fishes based on RNA-seq data originating from different muscle types (ventricle, hypaxial fast muscle, and ‘muscle’ respectively) depending on the species. It is important to note that such tissue independent changes in gene expression profiles are not uncommon (Stevens *et al*., 2013, Anastasiadi *et al*., 2018), especially when the stressor is introduced early on in development, prior to the specialisation of many tissues.

Changes in transcriptional entropy, of both the pairwise and higher order interaction networks, demonstrate a long-term effect of developmental temperature on the co-ordination of gene networks, which may contribute to the changed response of the transcriptome to environmental challenges in later-life (Figure 5). Indeed, our zebrafish re-analysis showed that the muscle tissue of fish reared in warmer conditions displays a greater number and magnitude of gene-expression changes than control animals following later-life cold-acclimation, as is noted in the original analysis by Scott and Johnston (2012). However, we also demonstrate that the entropy of zebrafish genes within the pairwise interaction networks is positively associated with the likelihood that the gene’s expression changes in response to the cold acclimation (Figure 5), but only in fish from the developmental warming background. These data suggest that developmental warming increases the thermal sensitivity of the transcriptome in later-life via changes in transcriptomic network organisation, but that transcriptome organisation is less important in the thermal acclimation process of fish from a ‘control’ background. Such complex interactions between developmental temperature and later-life plasticity, mediated through changes in regulatory mechanisms, have been demonstrated in previous studies (Loughland *et al*., 2021). Whilst this conclusion is tentative, we believe that the consistent changes in network entropy and their correlation with later-life gene expression changes following thermal acclimation warrant future investigation. Whilst the r^2^ of the logistic regressions were small, suggesting that the effect is both small and noisy at the level of individual genes, the cumulative effects across the whole transcriptome may be marked. Thus, the thermal environment experienced during embryogenesis is predicted to influence an individual’s responsiveness to future temperature challenges via changes in the entropy of the pairwise gene interaction networks. Future research is needed to further test this association and to discern the mechanisms facilitating the temperature-induced changes in transcriptomic organisation.

Recent studies have suggested that the developmental environment influences the capacity to acclimate to environmental changes in later-life (Beaman *et al*., 2016). For example, intertidal copepods (*Tigriopus californicus*) that undergo embryogenesis at 25 ºC have the capacity to raise their critical thermal maximum as adults through temperature acclimation (Healy *et al*., 2019).

However, embryos of the same species incubated at 20 ºC show no acclimation capacity in critical thermal maximum as adults (Healy *et al*., 2019). Given that phenotypic plasticity is likely facilitated by co-ordinated changes in gene expression, these developmentally plastic changes in later-life acclimation potential may relate to the differences in transcriptional organisation observed in our study. Further studies support the link between embryonic temperature and later-life acclimation capacity. Mosquitofish (*Gambusia holbrooki*) produce multiple generations per year, with those born in summer experiencing a warm but constant environment, and those born in spring experiencing cool, but steadily warming, conditions. Recent work has demonstrated that mosquitofish born in the more thermally variable spring environment have a greater capacity to acclimate their metabolic processes than mosquitofish from the more thermally stable summer conditions (Seebacher *et al*., 2014). Finally, studies of fruit flies (*Drosophila melanogaster*) have shown that heat tolerance is influenced not just by acclimation temperature, but by an interaction between acclimation temperature and embryonic temperature, further supporting the role of the early-life thermal environment in dictating an individual’s response to future temperature challenges (Willot *et al*., 2021).

Studies of developmental programming resulting from environmental challenges in fish and reptiles often show a protective phenotype in later-life, in contrast to the negative effects typically reported in mammals (Galli *et al*., 2021, Hellgren *et al*., 2021, Ruhr *et al*., 2019, Seeabacher *et al*., 2014). For example, work on the common snapping turtle (*Chelydra serpentina*) revealed that hypoxia exposure (50% air saturation) throughout embryogenesis improved the anoxia tolerance of the cardiomyocytes isolated from juveniles (Ruhr *et al*., 2019). Similarly, jacky dragons (*Amphibolurus muricatus*) raised with extended basking times (11 vs. 7 hours of daily heat lamp exposure) show a higher panting threshold than those from control conditions, implying a greater thermal tolerance (So and Schwanz, 2018). However, exposure to a stressor during embryogenesis does not always facilitate resilience to that stressor in adulthood. Cuban brown anole (*Anolis sagrei*) eggs incubated under cool, warm, and hot temperature fluctuations, and then raised in standard conditions after hatching, show no differences in thermal tolerance as adults (Gunderson *et al*., 2020). Thus, whilst developmental exposure to a stressor often influences the capacity to respond to that same stressor in later-life, it is not always the case.

One mechanism linking the developmental environment to later-life acclimation capacity is through the activity of DNA-methyltransferases (DNMTs) (Radford, 2018). DNA methylation by DNMTs can repress gene activity either directly, whereby the methylation prevents the interaction between a gene and its DNA binding proteins (Watt and Molloy, 1988), or indirectly, through recognition of the methyl cytosine by methyl cytosine binding proteins, and the consequent recruitment of transcriptional corepressors (Boyes and Bird, 1991, Klose and Bird, 2006). Changes in embryonic temperature have been shown to alter DNA methylation patterns in fish (Metzger and Schulte, 2017), with a recent study identifying DNMT3a as the mediator of developmental thermal plasticity in zebrafish (*Danio rerio*) (Loughland *et al*., 2021). Thus, by modulating DNMT3a’s activity, changes in embryonic temperature can have persistent effects on the plasticity of fishes (Loughland *et al*., 2021). Whilst the DEGs identified in our study show little overlap between the species (Figure 2), the observed changes in network entropy are present throughout the transcriptome and consistent across species. This transcriptome-wide remodelling suggests that a mechanism upstream of gene expression, such as epigenetic modifications, may be facilitating the developmentally programmed changes in gene network co-ordination. Indeed, epigenetic modifications are known to affect gene regulatory networks, for example, by altering the accessibility of chromatin (Yan *et al*., 2020). A promising avenue for future research is to investigate the role of DNMT genes in mediating the developmental plasticity of gene interaction networks.

Climate change is a major threat facing animals. As the world’s oceans and rivers continue to warm (Cai *et al*., 2014, Oliver *et al*., 2018), the physiological and population-level stresses exerted upon fishes will continue to grow. Embryogenesis is a sensitive period in many animal’s life histories, and the conditions experienced during embryonic development influence their growth and physiology (Galli *et al*., 2022, Dimitriadi *et al*., 2018, Scott and Johnston, 2012), as well as their capacity to respond to future stressors (Beaman *et al*., 2016, Seebacher *et al*., 2014, Healy *et al*., 2019, Scott and Johnston, 2012). Consequently, to predict and thus mitigate the consequences of anthropogenic warming, it is crucial that we begin to understand the mechanisms by which the developmental environment influences an organisms’ physiology and capacity to respond to future environmental challenges.

## Acknowledgements

We thank Vicky Taylor and the aquarist team in the Biological Services Facility at the University of Manchester for their assistance in animal husbandry. We thank Mar Pineda and Pierre Delaroche for their dissection skills. We thank Dr William Joyce, Dr Jade Taylor, and Professor David Eisner for thoughtful discussion of the project. We thank the aquarist team of the Ozeaneum, Stralsund for shark breeding. We thank Milton Tan and Lafage for their stickleback and seabass silhouettes shared, unchanged, through PhyloPic under the license: <u>Creative Commons — Attribution 3.0 Unported — CC BY 3.0</u>. Finally, we acknowledge the assistance given by Research IT and the use of the Computational Shared Facility at The University of Manchester.

## Author Contributions

D.M.R co-ordinated the experiments, analysed the data, and drafted the manuscript. T.G contributed to the code and analytical method development. S.A.H and A.V performed the microsatellite analysis, and S.A.H identified the individual animals. B.G and T.M supplied *Scyliorhinus canicula* eggs. B.G, T.M, and A.V reviewed the manuscript. H.A.S, P.C, and A.S conceived the study, secured funding, and reviewed and revised the manuscript. All authors contributed to the manuscript and gave their approval for publication.

## Data Availability

Novel sequencing data will be available upon publication through NCBI’s Gene Expression Omnibus (GEO, Edgar *et al*., 2002) at the GEO accession number GSE189976. The re-analysed data are available online through Metzger and Schulte 2018, Scott and Johnston 2012, and Anastasiadi *et al*. 2021.

**Supplementary Figure 1:**
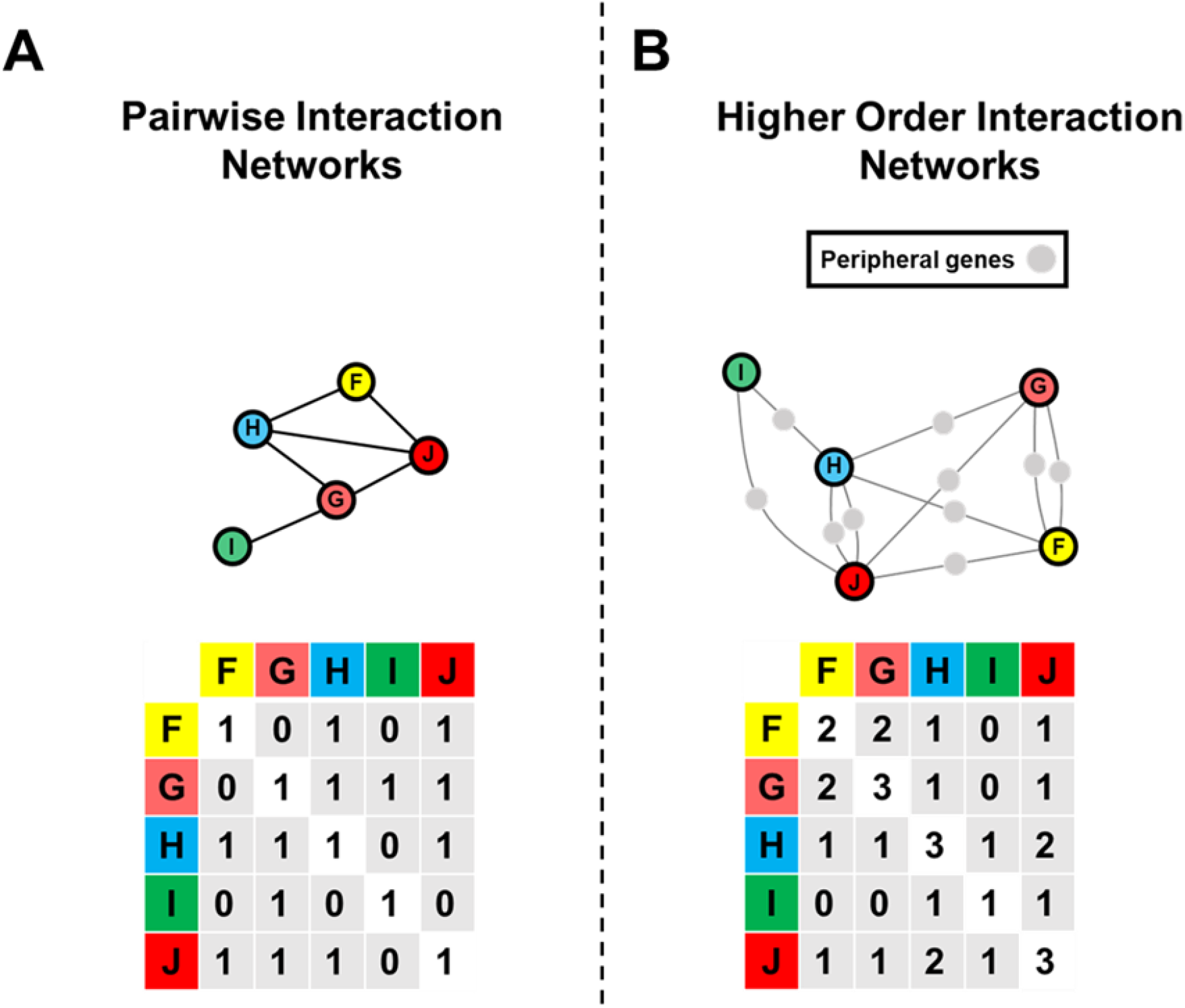
an example **A)** pairwise gene interaction network. Correlations between pairs of focal genes are shown by a solid black line and denoted by a 1 in the associated matrix. Pairwise gene network entropy is calculated from the pairwise interaction matrix and represents the network’s information content. **B)** the higher order interactions of the same example network. Focal genes (the two being assessed for their interaction) are associated via their shared connections to peripheral (not in the focal network) genes, denoted by the grey lines. An interaction between two focal genes can occur via one (gene H – gene I) or multiple (gene H – gene J) shared connections to the peripheral genes. The adjacency matrix denotes the number of shared connections to peripheral genes that each pair of focal genes has. Higher order network entropy is calculated from this adjacency matrix and represents the network’s information content.

## References

Anastasiadi, D., Diaz, N. and Piferrer, F. (2017) ‘Small ocean temperature increases elicit stage-dependent changes in DNA methylation and gene expression in a fish, the European sea bass’, Scientific Reports, 7.

Anastasiadi, D., Esteve-Codina, A. and Piferrer, F. (2018) ‘Consistent inverse correlation between DNA methylation of the first intron and gene expression across tissues and species’, Epigenetics & Chromatin, 11.

Anastasiadi, D., Shao, C. W., Chen, S. L. and Piferrer, F. (2021) ‘Footprints of global change in marine life: Inferring past environment based on DNA methylation and gene expression marks’, Molecular Ecology, 30(3), pp. 747–760.

Banerji, C. R. S., Miranda-Saavedra, D., Severini, S., Widschwendter, M., Enver, T., Zhou, J. X. and Teschendorff, A. E. (2013) ‘Cellular network entropy as the energy potential in Waddington’s differentiation landscape’, Scientific Reports, 3.

Bateson, P., Gluckman, P. and Hanson, M. (2014) ‘The biology of developmental plasticity and the Predictive Adaptive Response hypothesis’, Journal of Physiology-London, 592(11), pp. 2357–2368.

Battiston, F., Cencetti, G., Iacopini, I., Latora, V., Lucas, M., Patania, A., Young, J. G. and Petri, G. (2020) ‘Networks beyond pairwise interactions: Structure and dynamics’, Physics Reports-Review Section of Physics Letters, 874, pp. 1–92.

Beaman, J. E., White, C. R. and Seebacher, F. (2016) ‘Evolution of Plasticity: Mechanistic Link between Development and Reversible Acclimation’, Trends in Ecology & Evolution, 31(3), pp. 237–249.

Bolger, A. M., Lohse, M. and Usadel, B. (2014) ‘Trimmomatic: a flexible trimmer for Illumina sequence data’, Bioinformatics, 30(15), pp. 2114–2120.

Boyes, J. and Bird, A. (1991) ‘DNA methylation inhibits transcription indirectly via a methyl-cpg binding-protein’, Cell, 64(6), pp. 1123–1134.

Bray, N. L., Pimentel, H., Melsted, P. and Pachter, L. (2016) ‘Near-optimal probabilistic RNA-seq quantification’, Nature Biotechnology, 34(5), pp. 525–527.

Cai, O.R., Poloczanska, E.S., Brewer, P.G., Sundby, S., Hilmi, K., Fabry, V.J., Jung, S. (2014). ‘The Ocean In Climate: Change 2014: Impacts, Adaptation, and Vulnerability. Part B: Regional Aspects. Contribution of Working Group II to the Fifth Assessment Report of the Intergovernmental Panel on Climate Change [Barros, V.R., C.B. Field, D.J. Dokken, M.D. Mastrandrea, K.J. Mach, T.E. Bilir, M. Chatterjee, K.L. Ebi, Y.O. Estrada, R.C. Genova, B. Girma, E.S. Kissel, A.N. Levy, S. MacCracken, P.R. Mastrandrea, and L.L. White (eds.)]’. Cambridge University Press, Cambridge, United Kingdom and New York, NY, USA, 1655–1731.

Dimitriadi, A., Beis, D., Arvanitidis, C., Adriaens, D. and Koumoundouros, G. (2018) ‘Developmental temperature has persistent, sexually dimorphic effects on zebrafish cardiac anatomy’, Scientific Reports, 8.

Edgar, R., Domrachev, M. and Lash, A. E. (2002) ‘Gene Expression Omnibus: NCBI gene expression and hybridization array data repository’, Nucleic Acids Research, 30(1), pp. 207–210.

Fellous, A., Favrel, P. and Riviere, G. (2015) ‘Temperature influences histone methylation and mRNA expression of the Jmj-C histone-demethylase orthologues during the early development of the oyster Crassostrea gigas’, Marine Genomics, 19, pp. 23–30.

Galli, G. L. J., Ruhr, I. M., Crossley, J. and Crossley, D. A. (2021) ‘The Long-Term Effects of Developmental Hypoxia on Cardiac Mitochondrial Function in Snapping Turtles’, Frontiers in Physiology, 12.

Galli, G.L., Lock, M.C., Smith, K.L., Giussani, D.A. and Crossley 2nd, D.A. (2023). Effects of developmental hypoxia on the vertebrate cardiovascular system. Physiology, 38(2), pp.53–62.

Grabherr, M. G., Haas, B. J., Yassour, M., Levin, J. Z., Thompson, D. A., Amit, I., Adiconis, X., Fan, L., Raychowdhury, R., Zeng, Q. D., Chen, Z. H., Mauceli, E., Hacohen, N., Gnirke, A., Rhind, N., di Palma, F., Birren, B. W., Nusbaum, C., Lindblad-Toh, K., Friedman, N. and Regev, A. (2011) ‘Full-length transcriptome assembly from RNA-Seq data without a reference genome’, Nature Biotechnology, 29(7), pp. 644–U130.

Griffiths, A. M., Casane, D., McHugh, M., Wearmouth, V. J., Sims, D. W. and Genner, M. J. (2011) ‘Characterisation of polymorphic microsatellite loci in the small-spotted catshark (Scyliorhinus canicula L.)’, Conservation Genetics Resources, 3(4), pp. 705–709.

Gunderson, A.R., Fargevieille, A. and Warner, D.A. (2020). Egg incubation temperature does not influence adult heat tolerance in the lizard Anolis sagrei. Biology Letters, 16(1), p.20190716.

Haas, B. J., Papanicolaou, A., Yassour, M., Grabherr, M., Blood, P. D., Bowden, J., Couger, M. B., Eccles, D., Li, B., Lieber, M., MacManes, M. D., Ott, M., Orvis, J., Pochet, N., Strozzi, F., Weeks, N., Westerman, R., William, T., Dewey, C. N., Henschel, R., Leduc, R. D., Friedman, N. and Regev, A. (2013) ‘De novo transcript sequence reconstruction from RNA-seq using the Trinity platform for reference generation and analysis’, Nature Protocols, 8(8), pp. 1494–1512.

Healy, T. M., Bock, A. K. and Burton, R. S. (2019) ‘Variation in developmental temperature alters adulthood plasticity of thermal tolerance in Tigriopus californicus’, Journal of Experimental Biology, 222(22).

Hellgren, K. T., Premanandhan, H., Quinn, C. J., Trafford, A. W. and Galli, G. L. J. (2021) ‘Sex-dependent effects of developmental hypoxia on cardiac mitochondria from adult murine offspring’, Free Radical Biology and Medicine, 162, pp. 490–499.

Hook, S.A., McMurray, C., Ripley, D.M., Allen, N., Moritz, T., Grunow, B. and Shiels, H.A. (2019a). Recognition software successfully aids the identification of individual small-spotted catsharks Scyliorhinus canicula during their first year of life. Journal of Fish Biology, 95(6), pp.1465–1470.

Hook, S.A., Musa, S.M., Ripley, D.M., Hibbitt, J.D., Grunow, B., Moritz, T. and Shiels, H.A. (2019b). Twins! Microsatellite analysis of two embryos within one egg case in oviparous elasmobranchs. Plos one, 14(12), p.e0224397.

Iacono, G., Massoni-Badosa, R. and Heyn, H. (2019) ‘Single-cell transcriptomics unveils gene regulatory network plasticity’, Genome Biology, 20.

Klose, R. J. and Bird, A. P. (2006) ‘Genomic DNA methylation: the mark and its mediators’, Trends in Biochemical Sciences, 31(2), pp. 89–97.

Kopylova, E., Noe, L. and Touzet, H. (2012) ‘SortMeRNA: fast and accurate filtering of ribosomal RNAs in metatranscriptomic data’, Bioinformatics, 28(24), pp. 3211–3217.

Lesnoff, M., Lancelot, R. (2012). aod: Analysis of Overdispersed Data. R package version 1.3.2.

Loughland, I., Little, A. and Seebacher, F. (2021) ‘DNA methyltransferase 3a mediates developmental thermal plasticity’, Bmc Biology, 19(1).

Malecki, M. T., Jhala, U. S., Antonellis, A., Fields, L., Doria, A., Orban, T., Saad, M., Warram, J. H., Montminy, M. and Krolewski, A. S. (1999) ‘Mutations in NEUROD1 are associated with the development of type 2 diabetes mellitus’, Nature Genetics, 23(3), pp. 323–328.

Marshall, T. C., Slate, J., Kruuk, L. E. B. and Pemberton, J. M. (1998) ‘Statistical confidence for likelihood-based paternity inference in natural populations’, Molecular Ecology, 7(5), pp. 639–655.

McCarthy, D. J., Chen, Y. S. and Smyth, G. K. (2012) ‘Differential expression analysis of multifactor RNA-Seq experiments with respect to biological variation’, Nucleic Acids Research, 40(10), pp. 4288–4297.

McKinnon, J. S. and Rundle, H. D. (2002) ‘Speciation in nature: the threespine stickleback model systems’, Trends in Ecology & Evolution, 17(10), pp. 480–488.

Metzger, D. C. H. and Schulte, P. M. (2017) ‘Persistent and plastic effects of temperature on DNA methylation across the genome of threespine stickleback (Gasterosteus aculeatus)’, Proceedings of the Royal Society B-Biological Sciences, 284(1864).

Metzger, D. C. H. and Schulte, P. M. (2018) ‘Similarities in temperature-dependent gene expression plasticity across timescales in threespine stickleback (Gasterosteus aculeatus)’, Molecular Ecology, 27(10), pp. 2381–2396.

Murgas, K.A., Saucan, E. and Sandhu, R. (2022). Hypergraph geometry reflects higher-order dynamics in protein interaction networks. Scientific Reports, 12(1), p.20879.

Musa, S. M., Czachur, M. V. and Shiels, H. A. (2018) ‘Oviparous elasmobranch development inside the egg case in 7 key stages’, Plos One, 13(11).

Ogata, N., Kozaki, T., Yokoyama, T., Hata, T. and Iwabuchi, K. (2015) ‘Comparison between the Amount of Environmental Change and the Amount of Transcriptome Change’, Plos One, 10(12).

Oliver, E. C. J., Donat, M. G., Burrows, M. T., Moore, P. J., Smale, D. A., Alexander, L. V., Benthuysen, J. A., Feng, M., Sen Gupta, A., Hobday, A. J., Holbrook, N. J., PerkinsKirkpatrick, S. E., Scannell, H. A., Straub, S. C. & Wernberg, T. (2018). ‘Longer and more frequent marine heatwaves over the past century’, Nature Communications, 9.

Park, Y., Lim, S., Nam, J. W. and Kim, S. (2016) ‘Measuring intratumor heterogeneity by network entropy using RNA-seq data’, Scientific Reports, 6.

Pegado, M. R., Santos, C. P., Raffoul, D., Konieczna, M., Sampaio, E., Maulvault, A. L., Diniz, M. and Rosa, R. (2020) ‘Impact of a simulated marine heatwave in the hematological profile of a temperate shark (Scyliorhinus canicula)’, Ecological Indicators, 114.

Peiris, H., Raghupathi, R., Jessup, C. F., Zanin, M. P., Mohanasundaram, D., Mackenzie, K. D., Chataway, T., Clarke, J. N., Brealey, J., Coates, P. T., Pritchard, M. A. and Keating, D. J. (2012) ‘Increased Expression of the Glucose-Responsive Gene, RCAN1, Causes Hypoinsulinemia, beta-Cell Dysfunction, and Diabetes’, Endocrinology, 153(11), pp. 5212–5221.

Pörtner, H. O. and Knust, R. (2007) ‘Climate change affects marine fishes through the oxygen limitation of thermal tolerance’, Science, 315(5808), pp. 95–97.

R Core Team (2019). R: A language and environment for statistical computing. R Foundation for Statistical Computing, Vienna, Austria. URL https://www.R-project.org/.

Radford, E. J. (2018) ‘Exploring the extent and scope of epigenetic inheritance’, Nature Reviews Endocrinology, 14(6), pp. 345–355.

Riddell, E. A., Roback, E. Y., Wells, C. E., Zamudio, K. R. and Sears, M. W. (2019) ‘Thermal cues drive plasticity of desiccation resistance in montane salamanders with implications for climate change’, Nature Communications, 10.

Robinson, M. D., McCarthy, D. J. and Smyth, G. K. (2010) ‘edgeR: a Bioconductor package for differential expression analysis of digital gene expression data’, Bioinformatics, 26(1), pp. 139–140.

Rohart, F., Gautier, B., Singh, A. and Le Cao, K. A. (2017) ‘mixOmics: An R package for ‘omics feature selection and multiple data integration’, Plos Computational Biology, 13(11).

Ruane, P. T., Garner, T., Parsons, L., Babbington, P. A., Wangsaputra, I., Kimber, S. J., Stevens, A., Westwood, M., Brison, D. R. and Aplin, J. D. (2022) ‘Trophectoderm differentiation to invasive syncytiotrophoblast is promoted by endometrial epithelial cells during human embryo implantation’, Human Reproduction, 37(4), pp. 777–792.

Ruhr, I. M., McCourty, H., Bajjig, A., Crossley, D. A., Shiels, H. A. and Galli, G. L. J. (2019) ‘Developmental plasticity of cardiac anoxia-tolerance in juvenile common snapping turtles (Chelydra serpentina)’, Proceedings of the Royal Society B-Biological Sciences, 286(1905).

Ruhr, I., Bierstedt, J., Rhen, T., Das, D., Singh, S. K., Miller, S., Crossley, D. A. and Galli, G. L. J. (2021) ‘Developmental programming of DNA methylation and gene expression patterns is associated with extreme cardiovascular tolerance to anoxia in the common snapping turtle’, Epigenetics & Chromatin, 14(1).

Sanchez, A. (2019) ‘Defining Higher-Order Interactions in Synthetic Ecology: Lessons from Physics and Quantitative Genetics’, Cell Systems, 9(6), pp. 519–520.

Schaefer, J. and Ryan, A. (2006) ‘Developmental plasticity in the thermal tolerance of zebrafish Danio rerio’, Journal of Fish Biology, 69(3), pp. 722–734.

Schmieder, R. and Edwards, R. (2011) ‘Fast Identification and Removal of Sequence Contamination from Genomic and Metagenomic Datasets’, Plos One, 6(3).

Schnurr, M. E., Yin, Y. and Scott, G. R. (2014) ‘Temperature during embryonic development has persistent effects on metabolic enzymes in the muscle of zebrafish’, Journal of Experimental Biology, 217(8), pp. 1370–1380.

Scott, G. R. and Johnston, I. A. (2012) ‘Temperature during embryonic development has persistent effects on thermal acclimation capacity in zebrafish’, Proceedings of the National Academy of Sciences of the United States of America, 109(35), pp. 14247–14252.

Seebacher, F., Beaman, J. and Little, A. G. (2014) ‘Regulation of thermal acclimation varies between generations of the short-lived mosquitofish that developed in different environmental conditions’, Functional Ecology, 28(1), pp. 137–148.

Shannon, C. E. (1948). ‘A mathematical theory of communication’, Bell System Technical Journal, 27(3), pp. 379–423.

Simao, F. A., Waterhouse, R. M., Ioannidis, P., Kriventseva, E. V. and Zdobnov, E. M. (2015) ‘BUSCO: assessing genome assembly and annotation completeness with single-copy orthologs’, Bioinformatics, 31(19), pp. 3210–3212.

So, C. K. J. and Schwanz, L. E. (2018) ‘Thermal plasticity due to parental and early-life environments in the jacky dragon (Amphibolurus muricatus)’, Journal of Experimental Zoology Part a-Ecological and Integrative Physiology, 329(6-7), pp. 308–316.

Stevens, A., Hanson, D., Whatmore, A., Destenaves, B., Chatelain, P. and Clayton, P. (2013) ‘Human growth is associated with distinct patterns of gene expression in evolutionarily conserved networks’, Bmc Genomics, 14.

Teschendorff, A. E. and Enver, T. (2017) ‘Single-cell entropy for accurate estimation of differentiation potency from a cell’s transcriptome’, Nature Communications, 8.

Torson, A. S., Dong, Y. W. and Sinclair, B. J. (2020) ‘Help, there are ‘omics’ in my comparative physiology!’, Journal of Experimental Biology, 223(24).

Van Oosterhout, C., Hutchinson, W. F., Wills, D. P. M. and Shipley, P. (2004) ‘MICRO-CHECKER: software for identifying and correcting genotyping errors in microsatellite data’, Molecular Ecology Notes, 4(3), pp. 535–538.

Watt, F. and Molloy, P. L. (1988) ‘Cytosine methylation prevents binding to DNA of a hela-cell transcription factor required for optimal expression of the adenovirus major late promoter’, Genes & Development, 2(9), pp. 1136–1143.

Willot, Q., Ben, L. and Terblanche, J. S. (2021) ‘Interactions between developmental and adult acclimation have distinct consequences for heat tolerance and heat stress recovery’, Journal of Experimental Biology, 224(16).

Yan, F., Powell, D. R., Curtis, D. J. and Wong, N. C. (2020) ‘From reads to insight: a hitchhiker’s guide to ATAC-seq data analysis’, Genome Biology, 21(1).

Zhang, J. T. D., Hatje, K., Sturm, G., Broger, C., Ebeling, M., Burtin, M., Terzi, F., Pomposiello, S. I. and Badi, L. (2017) ‘Detect tissue heterogeneity in gene expression data with BioQC’, Bmc Genomics, 18.

Zhu, Z. Y., Surujon, D., Ortiz-Marquez, J. C., Huo, W. W., Isberg, R. R., Bento, J. and van Opijnen, T. (2020) ‘Entropy of a bacterial stress response is a generalizable predictor for fitness and antibiotic sensitivity’, Nature Communications, 11(1).

